# Derivatives of the antimalarial drug mefloquine are broad spectrum antifungal molecules with activity against drug-resistant clinical isolates

**DOI:** 10.1101/851816

**Authors:** Marhiah C. Montoya, Sarah Beattie, Kathryn M. Alden, Damian J. Krysan

## Abstract

The antifungal pharmacopeia is critically small, particularly in light of the recent emergence of multi-drug-resistant pathogens such as *Candida auris*. Herein, we report that derivatives of the anti-malarial drug mefloquine have broad spectrum antifungal activity against pathogenic yeasts and molds. In addition, the mefloquine derivatives have activity against clinical isolates that are resistant to one or more of the three classes of drugs currently used to treat invasive fungal infections, indicating that they have a novel mechanism of action. Importantly, the in vitro toxicity profiles using human cell lines indicate that the mefloquine derivatives are very similar to the parent mefloquine despite being up to 64-fold more active against fungal cells. In addition to direct antifungal activity, sub-inhibitory concentrations of the mefloquine derivatives inhibit the expression of virulence traits including filamentation in *C. albicans* and capsule formation/melanization in *C. neoformans*. Mode/mechanism of action experiments indicate that the mefloquine derivatives interfere with both mitochondrial and vacuolar function as part of a multi-target mechanism of action. The broad-spectrum scope of activity, blood-brain-barrier penetration, and large number of previously synthesized analogs available combine to support the further optimization and development of the antifungal activity of this general class of drug-like molecules.

## INTRODUCTION

The need for novel classes of antifungal drugs has never been more pressing (1). The numbers of patients who are at risk for developing invasive fungal infections continues increase as therapies that modulate the immune system are introduced into clinical practice. In addition, antifungal drug resistance has been increasing with resistance to all classes of antifungal drugs now well-recognized in pathogenic molds and yeast. Indeed, the recent emergence of the multi-drug resistant fungus, *Candida auris*, emphasizes the impact of this trend (2). Because we only have three classes of antifungal drugs, as opposed to at least ten classes of antibacterials, the options for treating fungal infections are extremely limited (1). As such, the path toward an untreatable fungal infection is extremely short. Although recent progress has been made with novel antifungal compounds moving into early-phase clinical trials, the pace of antifungal drug discovery must increase to insure a robust pipeline of new drugs. Accordingly, novel approaches to identify antifungal drug candidates are needed (3).

In many areas of medicine, repurposing existing drugs to new indications has been pursued as an expedited approach to developing new therapies (4). This type of search most commonly begins with a high-throughput screen (HTS) of a library of FDA-approved drugs. We and others have identified antifungal small molecules using this approach and two repurposed drugs have been evaluated as adjuvants in the treatment of cryptococcal meningoencephalitis (5, 6). Campaigns such as these have almost exclusively focused on identifying drugs that are candidates for direct translation to clinical use. However, a second type of repurposing approach involves the use of the identified molecule as a starting point for further optimization of the new activity. This has been termed <u>s</u>elective <u>o</u>ptimization of <u>s</u>ide <u>a</u>ctivities or SOSA (7).The rationale for this approach is based on the fact that it is quite difficult to predict whether or not a novel chemical scaffold will have drug-like properties or undesirable toxicities. Therefore, optimizing a desired activity within a drug-like scaffold with well-characterized pharmacologic properties and toxicities may lead to a clinically useful molecule more quickly than if development begins starting with a completely novel chemical entity. To our knowledge, this approach has not been widely applied to antifungal drug development.

We took a novel approach to identify a scaffold for SOSA-based optimization. Specifically, we performed manual literature searches for clinically used drugs that had been studied using genome wide, yeast-based chemical genetics experiments to characterize their mechanisms of action (8). Many of the drugs evaluated in the large-scale yeast chemical genetic screens had relatively low levels of antifungal activity that would not normally be sufficient to emerge from a high throughput screen. Consequently, this approach would allow us to access novel antifungal chemical space that could be amenable to further optimization. In addition, the chemical-genetic profiles of the drugs provide hypothesis-generating data regarding mechanism and mode of action.

Because most invasive fungal infections can affect the central nervous system in some manner (9), we narrowed our search to molecules that were known to cross the blood-brain-barrier. A drug that had been the focus of two yeast-based chemical genetics screens and fit our criteria was the antimalarial, mefloquine (MEF). MEF, previously WR 142490, is an orally available 4-quinoline-methanol (**Fig. 1A**) discovered by Experimental Therapeutics Division of the Walter Reed Army Institute of Research during the early 1970s to treat malaria and was FDA Approved in 1989 (10). An attractive feature of MEF was the availability of structural analogs of MEF through the NCI experimental therapeutics program.

**Figure 1.**
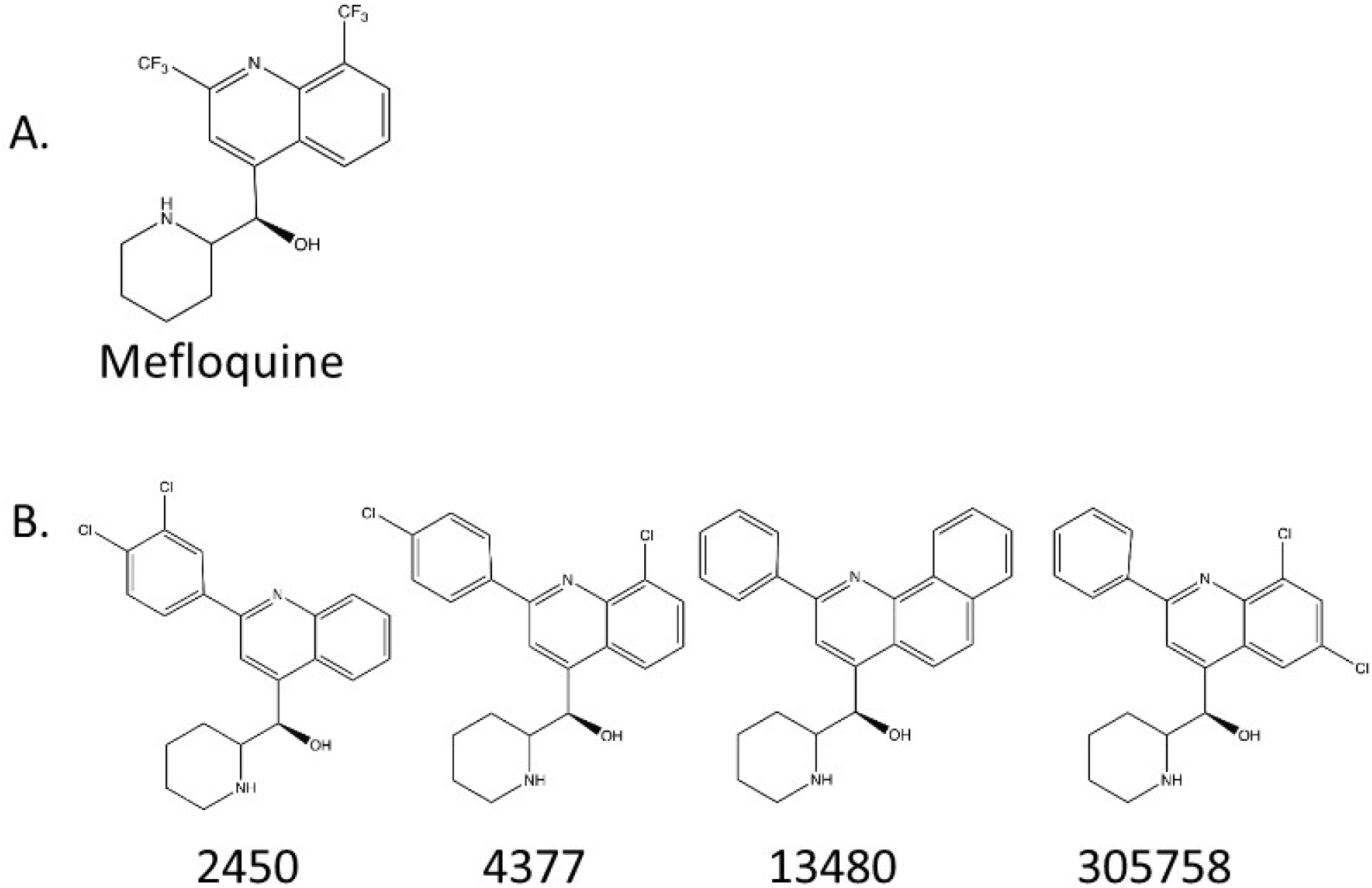
Chemical structures of mefloquine and mefloquine-derivatives. **A.** Mefloquine. **B.** Mefloquine derivatives obtained from National Cancer Institute Chemical Depository.

MEF has also been studied as a repurposing candidate against other human parasites including schistosomiasis, toxoplasmosis, and echinococcosis (11–17). MEF has also shown antibacterial activity against *Streptococcus pneumoniae* and *Mycobacterium tuberculosis* (18, 19). Additionally, repurposing efforts have found antiviral activity against Ebola, Dengue, and Zika virus (20, 21). Most importantly a small set of MEF-derivatives were shown to have activity against *Cryptococcus neoformans* and *Candida albicans* (22). Despite these promising findings, no additional characterization of the antifungal activity of the MEF-derivatives has been reported. Here, we show that MEF-derivatives have broad spectrum antifungal activity against both susceptible and drug-resistant fungi; show synergy with existing antifungal drugs; modulate the expression of virulence traits in both *C. albicans* and *C. neoformans*; and interfere with the functions and interactions of mitochondria and vacuoles in fungi.

## RESULTS

### Mefloquine derivatives have *in vitro* antifungal activity against a variety of human fungal pathogens

To explore the antifungal activity of the MEF scaffold, we first tested MEF against a set of reference strains of pathogenic fungi including *C. albicans*, *C. glabrata*, *C. auris*, *C. neoformans*, and *A. fumigatus* as well as the model yeast *S. cerevisiae* (**Table 1**). Consistent with previous reports, MEF has minimal antifungal activity, with *C. neoformans* being the most susceptible pathogenic species. As discussed above, Kunin et al. (22) had reported that some MEF-derivatives had improved antifungal activity relative to the parent drug. We, therefore, obtained 13 MEF-derivatives from the NCI chemical repository and screened them for antifungal activity. To our delight, four derivatives show improved antifungal activity relative to MEF (**Fig. 1B**, **Table 1**). The most active derivative is 4377 and has minimum inhibitory concentration (MIC) values below 4 µg/mL for all species tested; 4377 was not previously examined by Kunin et al. (22).

**Table 1.** Minimum Inhibitory Concentrations for Mefloquine Derivatives Against Susceptible Fungal Reference Strains

| Species | Strain | Minimum Inhibitory Concentration (µg/ml) |  |  |  |  |  |  |  | VOR |
| --- | --- | --- | --- | --- | --- | --- | --- | --- | --- | --- |
|  |  | MEF | 2450 | 4377 | 13480 | 305758 | AmB | FLU | CAS |  |
| *C. albicans* | SC5314 | >128 | 8 | 4 | 4-8 | 8 | 0.25 | 0.25 | 0.125-0.25 | - |
| *C. glabrata* | BG2 | 128 | 8 | 4 | 2-4 | 8 | 1-2 | 8 | 0.5 | - |
| *C. auris* | 0381 | 128 | 8 | 2 | 4 | 4 | 0.38<sup>a</sup> | 4<sup>a</sup> | 0.125<sup>a</sup> | - |
| *C. neoformans* | H99 | 32 | 4 | 1 | 1 | 2 | 0.5-1 | 2-4 | - | - |
| *S. cerevisiae* | BY4741 | >16 | 8 | 4 | >16 | 16 | - | >16 | 0.03 | - |
|  | BY4742 | >16 | 8 | 4 | 16 | 8 | - | >16 | 0.03 | - |
|  | BY4743 | >16 | 8 | 4 | 16 | 8 | - | >16 | 0.03 | - |
| *A. fumigatus* | AF293 | > 64 | 8 | 2 | 4 | 4 | - | - | - | 1 |
|  | CEA10 | >64 | 8 | 2 | 4 | 8 | - | - | - | 0.5 |
|  | SPF93 | >64 | 8 | 2 | 4 | 8 | - | - | - | 0.5 |

<sup>a</sup> MIC reported by Centers for Disease Control and Prevention

To confirm the antifungal activity of the MEF-derivatives, we tested them against a variety of isolates with varying susceptibilities to clinically used antifungal drugs including fluconazole- and echinocandin-resistant *Candida* spp. and voriconazole-resistant *A. fumigatus*. As shown in **Table 2**, the MICs for the MEF-derivatives did not vary more than 2-fold amongst this diverse set of clinical fungal isolates. Most importantly the MEF-derivatives were active against strains such as *C. albicans* strain TWO7343 with increased multi-drug efflux activity (23), indicating that the MEF-derivatives are not suceptibile to efflux pumps that are active against azoles. The MEF-derivatives are also uniformly active against *C. auris* isolates with decreased susceptibility to fluconazole (FLU), caspofungin (CAS), and amphotericin B (AmB) (**Table 2**). These data firmly establish this scaffold as having broad spectrum antifungal activity.

**Table 2.** Minimum Inhibitory Concentrations of Mefloquine Derivatives Against Antifungal-Resistant Clinical Isolates

| Species | Clinical Isolate ID | Minimum Inhibitory Concentration (µg/ml) |  |  |  |  |  |  |  |  |
| --- | --- | --- | --- | --- | --- | --- | --- | --- | --- | --- |
|  |  | MEF | 2450 | 4377 | 13480 | 305758 | AmB | FLU | CAS | VOR |
| *C. albicans* | 1010 | >128 | 8 | 4 | 8 | 8 | 0.5 | > 256 | 4 | - |
|  | O-788 | >128 | 8 | 4 | 8 | 8 | 0.5 | > 256 | 2 | - |
|  | TWO7243 #17 | >64 | 8 | 4 | 8 | 16 | 1 | >128 | 0.25 | - |
| *C. glabrata* | 1 | 128 | 8 | 2 | 2 | 4 | 0.5 | 2 | 2-4 | - |
|  | 102 | >64 | 8 | 4 | 2-4 | 8 | 0.5 | ≥64 | 64 | - |
|  | 999 | >128 | 8 | 4 | 4 | 16 | 0.5 | 128 | 64 | - |
|  | 4720 | 128 | 8-16 | 2-4 | 4 | 8 | 0.25-0.5 | 128 | 64 | - |
| *C. auris* | 0383 | 128 | 16 | 4 | 4 | 8 | 0.38<sup>a</sup> | 128<sup>a</sup> | 0.25<sup>a</sup> | - |
|  | 0385 | 128 | 8 | 4 | 4 | 8 | 0.5<sup>a</sup> | >256<sup>a</sup> | 0.5<sup>a</sup> | - |
|  | 0387 | 128 | 8 | 2 | 8 | 8 | 0.75<sup>a</sup> | 8<sup>a</sup> | 0.25<sup>a</sup> | - |
|  | 0388 | 128 | 8 | 4 | 8 | 8 | 1.5<sup>a</sup> | >256<sup>a</sup> | 1<sup>a</sup> | - |
|  | 0390 | 128 | 16 | 4 | 8 | 8 | 4<sup>a</sup> | >256<sup>a</sup> | 0.5<sup>a</sup> | - |
| *C. neoformans* | 101.14 | 64 | 4 | 2 | 2 | 4 | 1 | 16 | - | - |
|  | 103.98 | 64 | 4 | 2 | 2 | 4 | 1 | 4 | - | - |
|  | 111.00 | 32 | 2 | 1 | 1 | 2 | 1-2 | 2 | - | - |
|  | 118.00 | 64 | 4-8 | 2 | 2-4 | 4 | 0.5-1 | >16 | - | - |
|  | 120.04 | 32 | 4-8 | 2 | 2 | 4-8 | 0.5-1 | 4 | - | - |
|  | 127.01 | 32 | 4 | 1 | 2 | 2 | 1-2 | 1 | - | - |
|  | 138.97 | 32 | 4 | 1 | 1 | 2 | 1 | 4-8 | - | - |
|  | 153.02 | 32 | 4 | 1 | 2 | 2 | 2 | 4 | - | - |
|  | 158.03 | 32 | 4 | 1 | 2 | 2 | 2 | >16 | - | - |
|  | 226.03 | 64 | 4 | 2 | 2 | 4 | 2 | 16 | - | - |
| *A. fumigatus* | SPF98 | >64 | 8 | 2 | 4 | 8 | - | - | - | 4 |

746 <sup>a</sup> MIC reported by Centers for Disease Control and Prevention

Finally, we attempted to isolate strains that were resistant to the MEF-derivatives by serial passage of strains through increasing concentrations of the compounds. Although some initially resistant colonies were identified, none were stably resistant and all reverted to the parental susceptibility following passage on compound-free media. As such, the MEF-derivatives have low rates of sponateous or selected resistance development. Overall, these derivatives of MEF have broad spectrum activity against medically important fungi and are not affected by resistance mechanisms against currently used antifungal drugs.

### Combination of mefloquine or mefloquine derivatives are fungicidal

Combination antifungal therapy is of increasing interest as a means to modulate antifungal resistance and/or improve antifungal efficacy (24). To assess the interactions between currently used antifungal drugs and MEF-derivatives, we performed checkerboard microdilution assays and analyzed data using the fractional inhibitory concentration index (FICI). FICIs were determined for *C. albicans* (SC5314), *C. glabrata* (BG2), and *C. neoformans* (H99) and the *C. glabrata* (#102), a clinical isolate resistant to both FLU and CAS. We did not observe any synergistic interactions between the MEF-derivatives and either CAS or FLU; all were additive (0.5<FICI<1.0, **Table S1**). Interestingly, 4377 at ½ MIC reduced the concentration of CAS to inhibit growth by >64-fold in *C. glabrata* isolate #102. Although this is formally an additive interaction because it was only observed at ½ MIC of 4377, it indicates that the susceptibility of the strain to CAS is dramatically modulated by 4377.

A second type of clinically-relevant interaction between two drugs is the conversion of drugs that are fungistatic by themselves to a combination that is fungicidal. This type of interaction is particularly relevant to the treatment of cryptococcal meningitis because fungicidal activity has been shown to improve outcomes (25). Specifically, outcomes for patients treated with fungistatic FLU are much worse than patients treated with fungicidal AmB (26). Accordingly, the identification of molecules that lead to fungicidal combinations with FLU has been pursued as an approach to improving the treatment of cryptococcal meningitis (27). Therefore, we investigated the killing kinetics of MEF and MEF-derivatives combined with FLU against *C. neoformans* using time-kill analysis. As shown in **Fig. 2A**, the combination of fungistatic concentrations of MEF with FLU is fungicidal and sterilizes the culture. Similar behavior was observed with MEF-derivatives with all but 13480 showing >2 Log_10_ reductions in CFU by 24 hours (**Fig. 2B**). These data indicate that combinations of MEF-derivatives with FLU may lead to improved efficacy relative to FLU alone for *C. neoformans*.

**Figure 2.**
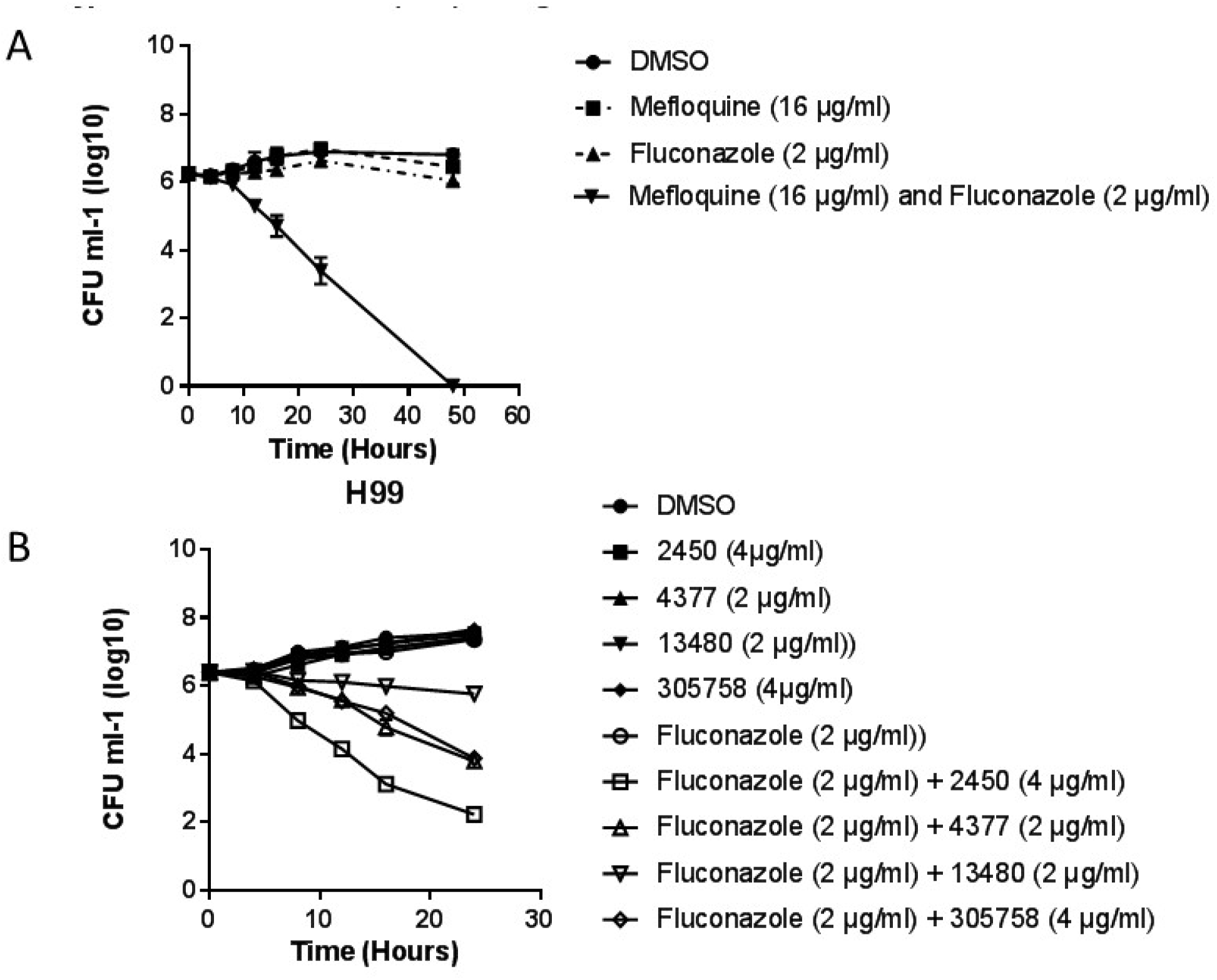
The combination of mefloquine and mefloquine-derivatives with fluconazole is fungicidal against *Cryptococcus neoformans*. **A.** The combination of fungistatic concentrations of mefloquine (1/4 MIC) and fluconazole (1/2 MIC) is fungicidal for strain H99 incubated in YPD at 37°C. **B**. All of the mefloquine derivatives except for 13480 showed similar fungicidal activity in combination with fluconazole under the conditions described above. Samples of each culture were plated on YPD at the indicated time points and the colony forming units/mL (CFU/mL) were quantified using serial dilutions after incubation at 30°C. The curves are single experiments representative of at least two independent experiments. Error bars indicate the standard deviation of two technical replicates.

### The in vitro toxicity of mefloquine derivatives against human cell lines is similar to mefloquine

Mefloquine is an FDA-approved drug for malaria and its toxicity has been characterized extensively. MEF is more toxic against immortalized human cell lines than it is against primary cell lines (28); however, these data provide a measure of the relative toxicity of new derivatives compared to the clinically used MEF. Therefore, we determined the LD_50_ for mefloquine and its derivatives against both HepG2 and A549 cells using metabolic activity assays. As shown in **Fig. 3A&B**, the IC_50_ for none of the derivatives differed from mefloquine by approximately two-fold. The antifungal activity for the derivatives improved by 8->64-fold. Therefore, the improved antifungal activity for the derivatives cannot be attributed to a general increase in eukaryotic toxicity. Although MEF has relatively poor antifungal activity, it achieves very high tissue concentrations near its MIC against organisms such as *Cryptococcus* (29). The relatively small changes in toxicity associated with the increased antifungal activity are very encouraging and suggest the scaffold may be amenable to *in vivo* therapy.

**Figure 3.**
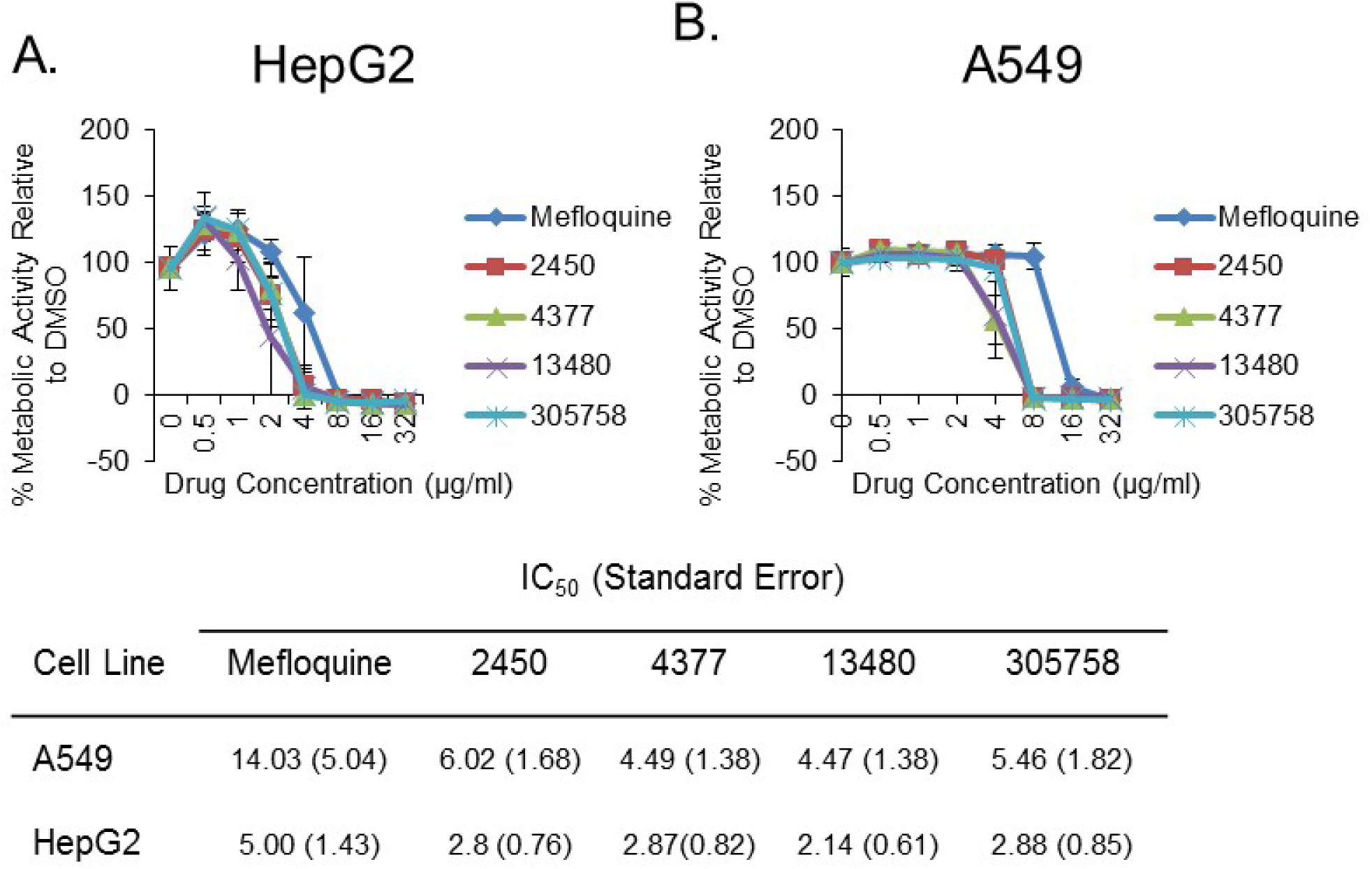
*In vitro* toxicity of mefloquine and mefloquine-derivatives against two cells lines are similar. The indicated cell lines (**A**: HepG2 and **B**: A549) were treated with DMSO or the indicated mefloquine derivative. The metabolic activity of the cells was determined by measuring the reduction of a metabolically active dye and compared to untreated controls over the indicated range of drug concentrations. The curves are representative of three independent experiments and the error bars indicate the standard deviation of three technical replicates.

### Mefloquine and mefloquine derivatives modulate fungal virulence factors

Fungal virulence traits have long been studied for their contribution to disease pathogenesis and, recently, interest in targeting these processes for the development of new therapies has increased (30). Among the *C. albicans* virulence traits, hyphal morphogenesis has been one of the most extensively studied both in terms of pathogenesis and molecular targeting (31). Indeed, another quinolone anti-malarial, quinacrine, inhibits *C. albicans* filamentation (32). We, therefore, examined the effect of MEF and its derivatives on *C. albicans* morphogenesis.

To investigate if MEF and the mefloquine derivatives affected *C. albicans* ability to form hyphae, the *C. albicans* reference strain SC5314 was exposed to sublethal concentrations of the molecules under conditions that induce filamentation (1% fetal bovine serum in YPD at 37°C). At concentrations of one-half MIC, MEF and its derivatives all suppress filamentation; almost all treated cells remain in yeast form but are still viable by propidium iodide (PI) staining (**Fig. 4A)**. MEF showed dose-dependent inhibition of filamentation (**Fig. 4B**), while the MEF-derivatives all completely inhibited filamentation at ½ MIC (**Fig. 4C**). Taken together with previously published quinacrine data, this indicates that the quinolone class appears to be a general inhibitor of *C. albicans* filamentation (33).

**Figure 4.**
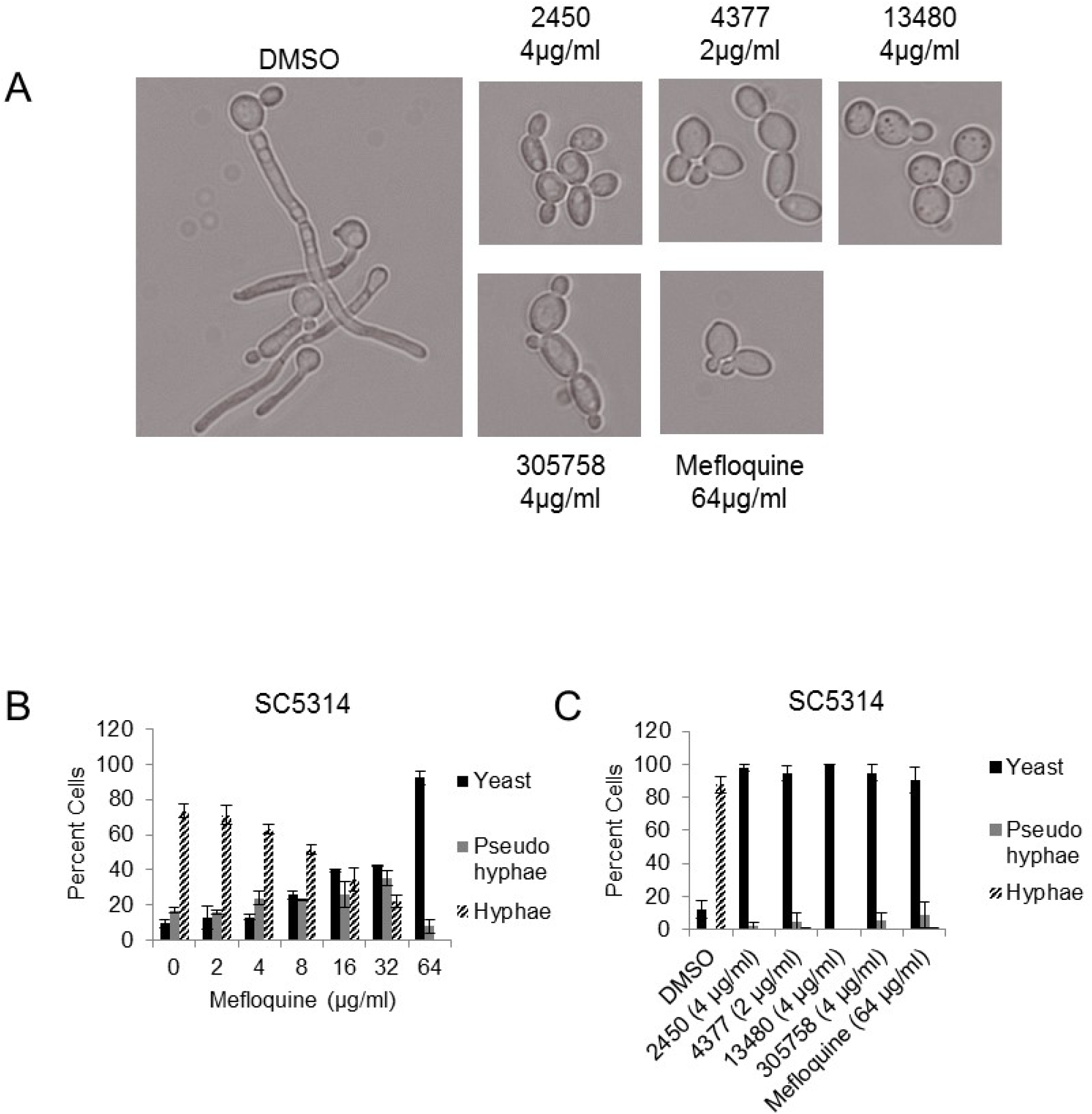
Mefloquine and mefloquine-derivatives inhibit *C. albicans* filamentation at sub-growth inhibitory concentrations. *C. albicans* strain SC5314 was inoculated into YPD supplemented with 1% fetal bovine serum at 37°C in the presence of either DMSO or the indicated concentration of a MEF derivative. After 2 hours, samples of the cultures were stained with propidium iodide, harvested, and examined by microscopy.

Two of the most important virulence characteristics of *C. neoformans* are polysaccharide capsule formation and melanization (34, 35). The polysaccharide capsule protects is unique among fungi and protects the cells against a variety of stress including host defense (34, 36). Importantly, *C. neoformans* strains that are not able to form capsule are avirulent in mouse models (37). Melanization is also thought to protect cells against oxidative stress such as macrophage generated reactive oxygen species. Recent patient-derived data have shown that treatment outcome correlates with extent of melanin formation (38). Thus, a drug that prevents capsule and melanin formation may be useful as an adjuvant, even if it is not sufficiently active to directly kill *C. neoformans*. (37).

*C. neoformans* H99 was exposed to sub-inhibitory concentrations of MEF and MEF-derivatives in capsule-inducing conditions and capsule formation was evaluated at 48 h by India ink staining. Treated cells were stained with PI before imaging to confirm that they were still viable. All MEF-derivatives caused a reduction in the number of cells with capsule and a decrease in size of capsule compared to DMSO (**Fig. 5A&B**). Similarly, the MEF-related compounds reduced melanization of *C. neoformans* cells incubated on L-DOPA plates (**Fig. 5C**). Taken together, these data indicate that MEF-derivatives modulate the expression of key virulence properties of pathogenic fungi and directly kill the organisms.

**Figure 5.**
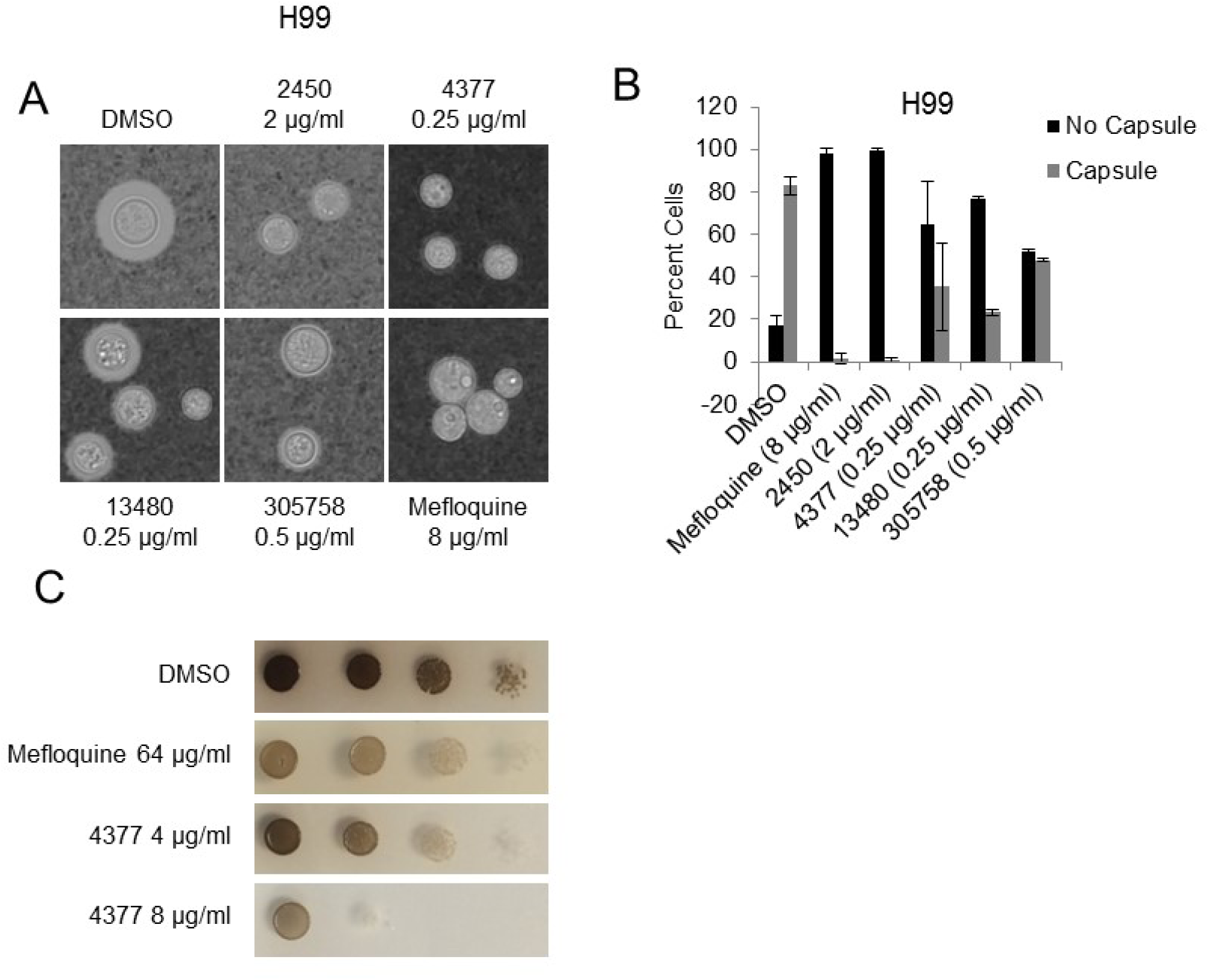
Mefloquine and mefloquine-derivatives suppress the expression of *C. neoformans* virulence traits. **A&B**. *Cryptococcus neoformans* strain H99 was incubated in DMEM at 37°C in 5% CO_2_ in the presence of DMSO or the indicated concentration of mefloquine derivative (1/2 to 1/4 MIC). The cells were harvested and stained with India Ink. The cells were photographed (**A**) and the number of cells with or without capsule counted (N > 100 cells per experiment). The bars in **B** indicate the mean of two or three independent experiments and the error bars are the standard deviations. **C**. A series of ten-fold dilutions of H99 was spotted on plates containing L-DOPA medium as well as the indicated concentration of mefloquine derivative, incubated at 37°C for 48 hr, and photographed. The photograph is representative of two independent replicates.

### Mode and mechanism of the antifungal activity of the mefloquine scaffold

As alluded to in the introduction, the mechanism of action for MEF as an anti-malarial drug has remained enigmatic despite its many years of use in the clinical setting. Recent proposals for its mechanism of action either as an antimalarial drug or as part of repurposing to other indications include: lysosomal/vacuole inhibitor (39), disruptor of mitochondrial proton motive force (40), protein translation inhibitor (41), and purine nucleoside phosphorylase (PNP) inhibitor (42). The latter two proposals involve specific target proteins, a subunit of 80S ribosome and the PNP enzyme. The yeast PNP enzyme is not essential and, thus, this target was eliminated from further consideration. We tested the effect of MEF and MEF-derivatives on yeast protein translation but found no evidence of inhibition (data not shown). Soon after we completed these experiments, Sheridan et al. reported that they also did not observed any effect of MEF on malarial cell protein translation (43).

MEF was identified as a potential adjuvant therapy for chronic myeloid leukemia and, as part of a multi-disciplinary effort to understand its mechanism on anti-cancer activity, a yeast-based chemical haploinsufficiency screen was performed to identify *S. cerevisiae* genes deletions that are hyper-susceptible to MEF (39). The set of genes that emerged from this screen was enriched for genes involved in yeast vacuole and protein secretion processes. Follow-up cell biological studies using human cells were consistent with this potential mechanism; however, no experiments involving yeast were reported.

To investigate the effect of MEF-derivatives on yeast vacuoles, log phase *S. cerevisiae* cells (BY4741) were exposed to sub-inhibitory concentrations of compound or DMSO for 2 h, stained with FM4-64, and imaged. All MEF-derivatives caused gross changes in vacuole morphology (**Fig. 6**); similar results were observed with *C. albicans* (data not shown). The changes begin with segmentation and invagination of a single vacuole and lead to many cells having an increased number of fragmented vacuoles. Many cells showed a single large vacuole surrounded by a varying number of smaller vacuoles that resembled class F vacuolar protein sorting mutants (44), which include *VPS1* and *VPS26* deletions. To directly compare phenotypes, we stained *vps1Δ S. cerevisiae* (BY4741 background) alongside DMSO or drug treated cells and indeed many cells showed vacuoles with morphology similar to *vps1Δ* (**Fig. 6B**).

**Figure 6.**
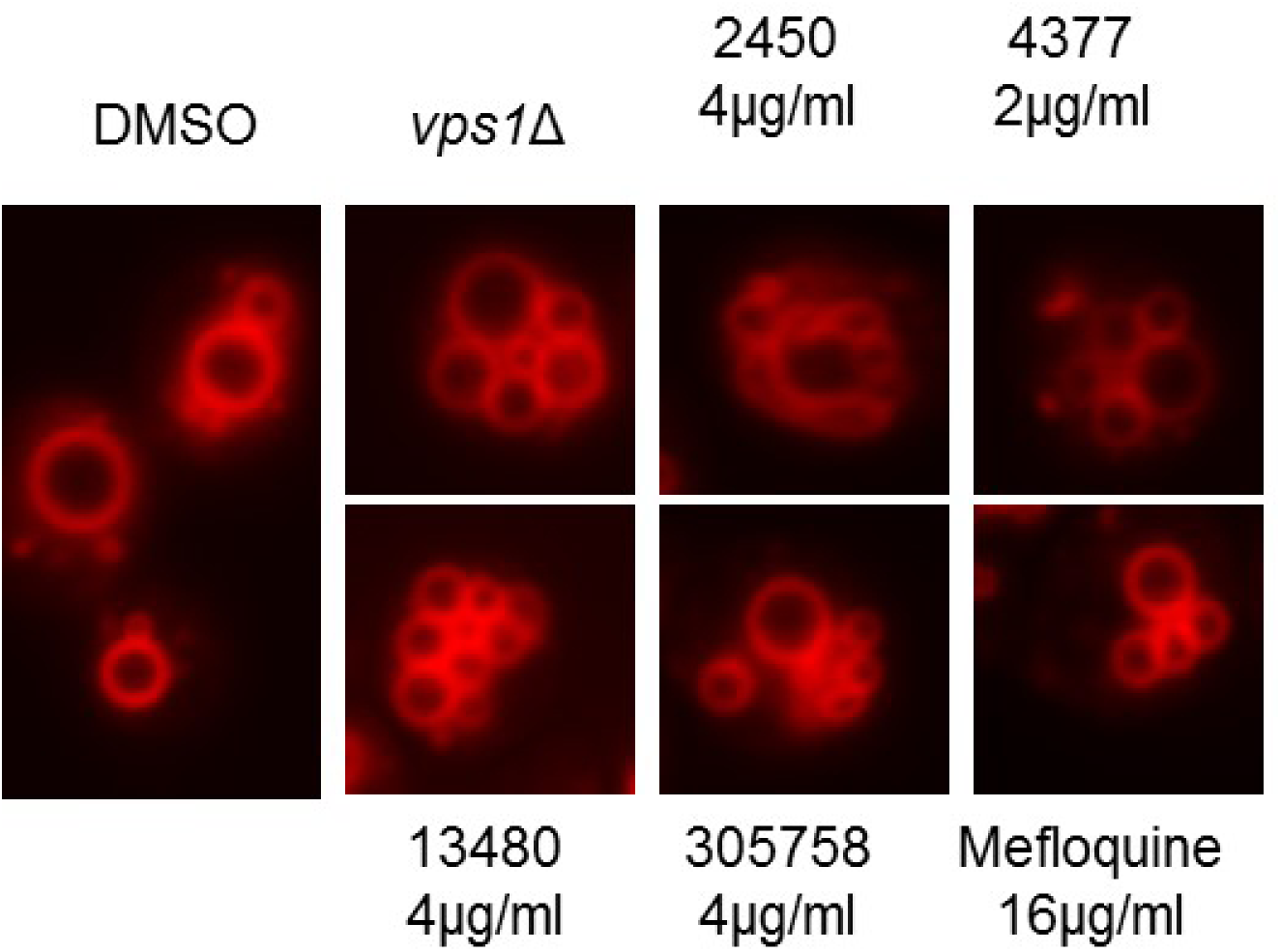
Mefloquine-derivatives disrupt vacuolar morphology. Exponential phase cultures of *S. cerevisiae* strain BY4741, or a derivative containing a *vps1*Δ mutation, were exposed to DMSO or the indicated drug concentrations in YPD at 30°C for 2 hours. The cells were harvested and stained with FM4-65 to visualize the vacuole and photographed. The vacuolar morphology of treated cultures is similar to the pattern reported for the *vps1*Δ strain. Propidium iodide straining of the same cultures showed that >95% of cells were viable at the time of processing.

Although the morphological effects of the MEF-derivatives were not as uniform as observed with *vps1*Δ, it seemed possible that Vps1 could be a target. Many *VPS* mutants mis-localize the vacuolar protein, Carboxypeptidase Y (CPY), to the extracellular space. However, we found that exposure of BY4741 to the MEF-derivatives did not lead to increased secretion of CPY by colony blot assays (data not shown). The *vps1*Δ mutant has a strong CPY secretion phenotype Although this assay is not as sensitive as radiolabeling assays, it seems likely that the MEF-derivatives do not have a profound effect on CPY mis-localization. Additionally, none of the MEF-derivatives inhibited recombinant Vps1 *in vitro* (data not shown). Therefore, while MEF and its antifungal derivatives clearly effect vacuole morphology, it seems unlikely that this effect is the sole mode of antifungal action.

### Mefloquine and mefloquine derivatives dissipate mitochondrial membrane potential and induce petite formation

MEF has been shown to interfere with mitochondrial function in a variety of eukaryotic systems. Feng et al. found that MEF was among a large set of clinically used drugs that dissipate the proton motive force across the mitochondrial membrane of eukaryotic cells, including *S. cerevisiae* (40). The anti-malarial activity of MEF has been attributed to mitochondrial dysregulation and production of reactive oxygen species (ROS) (45). A similar mitochondrial effect has been reported in cervical cancer cell lines and results in a decrease in ATP levels as well as increased cellular ROS that lead to cell death (46). To determine if MEF-derivatives affect the mitochondrial proton motive force in pathogenic yeast, log phase *C. albicans* cells (SC5314) were exposed cells to sub-inhibitory concentrations of drug or DMSO for 2 h, stained with MitoTracker Red CMXRos and Green, and imaged. MitoTracker Red uptake is dependent on the mitochondrial membrane potential while Green is not. As shown for 13480, MitoTracker Red uptake was nearly abolished while the morphology of the mitochondria was unchanged (**Fig. 7A**). The proton motive force was also decreased in *C. neoformans* (**Fig. 7B**). Additionally, similar results were obtained for 2450 (4 µg/ml), 4377 (2 µg/ml), 305758 (4 µg/ml), and MEF (64 µg/ml), indicating that these drugs dissipate the mitochondrial proton motive force as part of its antifungal activity (**Fig. 7C**).

**Figure 7.**
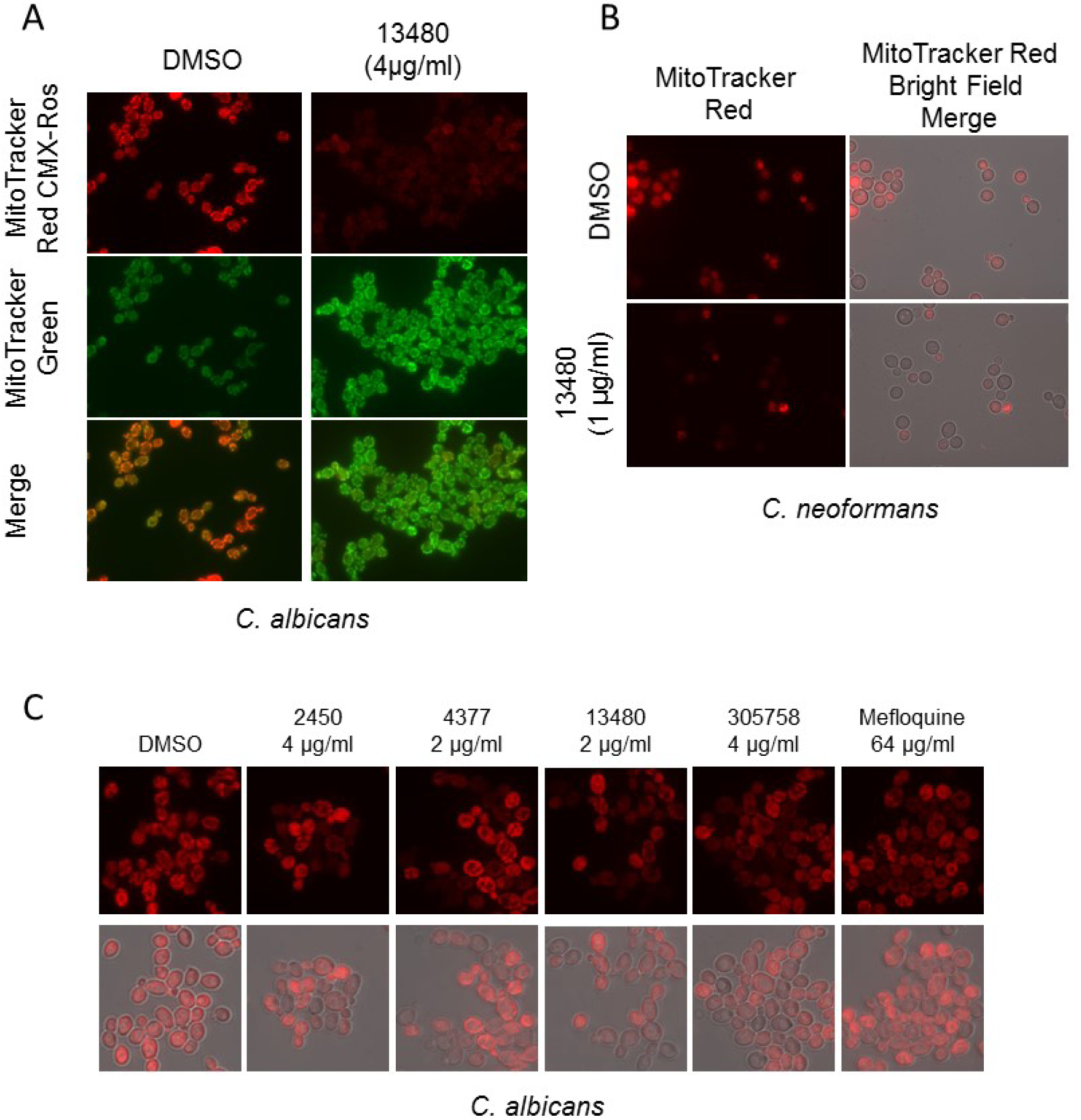
Mefloquine-derivatives dissipate proton motive force across the mitochondrial membrane. **A**. *C. albicans* SC5314 cells were treated with the indicated concentration of 13480 or DMSO for 2 hr in YPD at 37°C. The cells were harvested, stained with either Mitotracker Red or Mitotracker Green and imaged. Identical exposure settings were used for both samples. Propidium iodide staining indicated that >95% cells were viable at the time of imaging. **B**. *Cryptococcus neoformans* H99 were processed similarly but for Mitotracker Red staining only. **C**. Other mefloquine derivatives show decreased uptake of Mitotracker Red against *C. albicans*.

Two recently identified antifungal small molecules that directly target the mitochondria, T-2307 (47) and Inz-1 (48), have increased activity in non-fermentable carbon sources such as glycerol as compared to standard glucose media. If mitochondrial disruption is the process driving the majority of MEF-derivative antifungal activity, we reasoned that the MIC for the MEF-derivatives should be much lower in the presence of glycerol; however, the MICs of the MEF-derivatives were decreased modestly (e.g., 2-fold; data not shown) for some derivatives or were unchanged. In addition, we compared the activity of MEF-derivatives against *C. glabrata* KK2001 and a respiratory-deficient rho*^0^*-derivative of KK2001, which has no functioning mitochondrion. The MICs were identical for the parent KK2001 strain and the rho^0^ KK2001 strain for 2450 and 4377. If disruption of mitochondrial function was the sole activity of the MEF scaffold, then rho*^0^*-strains should be resistant. Therefore, these data further suggest that the mitochondrion is targeted by the MEF scaffold and that such activity is not solely responsible for its antifungal properties.

Interestingly, incubation of the parent KK2001 strain in the presence of MEF analog for 48 h led to some growth beyond the initial MIC observed at 24 h. To investigate the possible explanation for supra-MIC growth, the cells from wells containing 305758 (8 µg/ml) and DMSO only were plated in 10-fold dilutions on YPD plates and incubated at 30°C for 24 h. After 24 h, the cells treated with 305758 grew as a mixture of normal sized and petite colonies (**Fig. 8A**), while plates of the DMSO-exposed cells had norm al sized colonies only (data not shown). We suspected these cells may be respiratory-deficient, petite colonies. To confirm the petite phenotype, 15 petite colonies and 4 controls (2 DMSO only colonies and 2 normal sized colonies from 305758 8 µg/ml) were patched on YP+2% glycerol (YPG) plates and incubated at 30°C for 24 h. None of the 15 petite colonies grew on YPG plates while all control colonies showed the expected amount of growth (**Fig. 8B**). This observation suggests that MEF-derivatives also affect mitochondrial genome stability.

**Figure 8.**
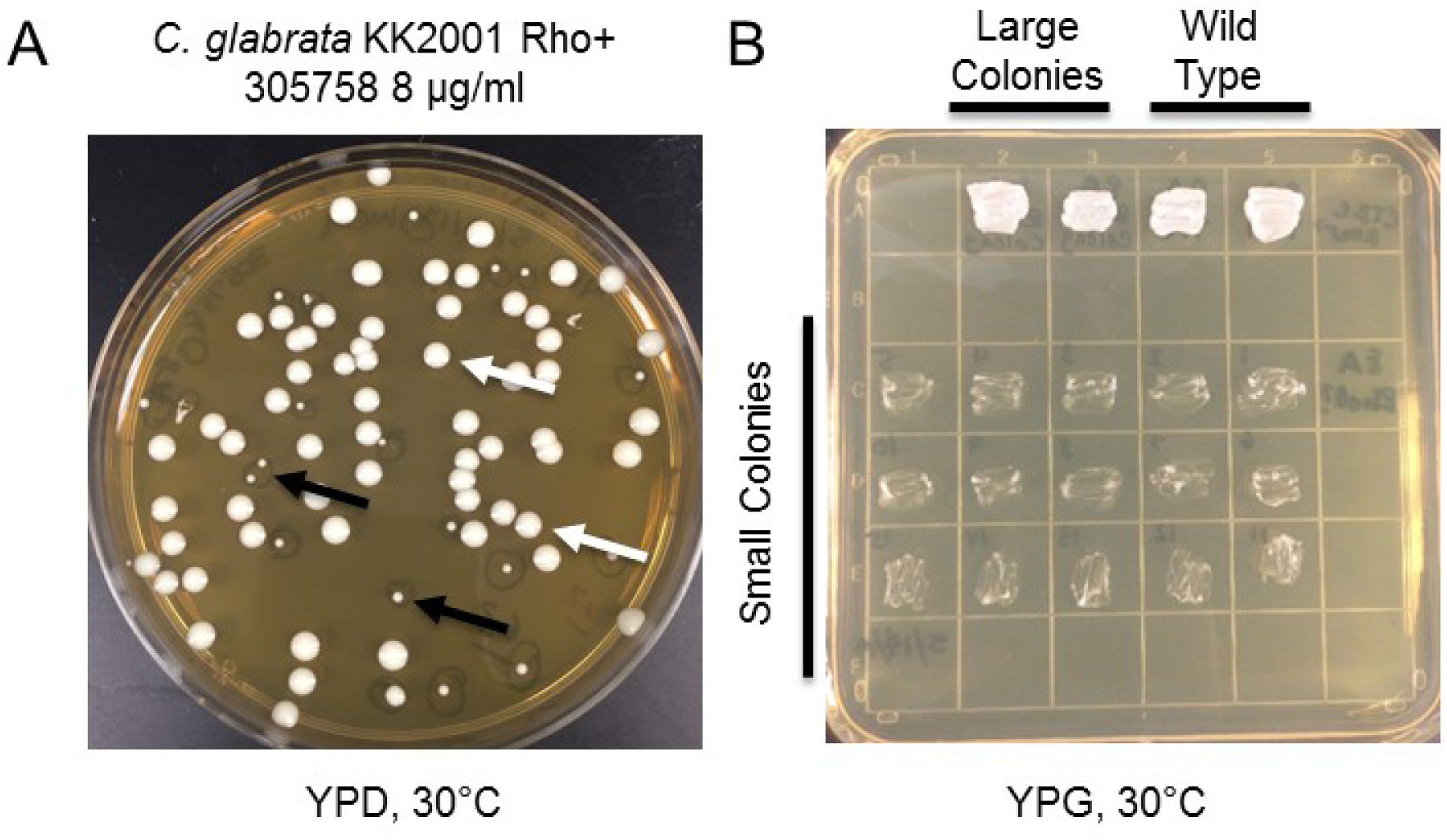
Mefloquine-derivatives induce mitochondrial DNA instability in *Candida glabrata*. **A.** *C. glabrata* strain KK2001 was incubated with 4377 (1/2 MIC) overnight in YPD medium. The surviving cells were plated on YPD, incubated at 30°C, and photographed. The black arrows indicate small or petite colonies while the white arrows indicate normal sized colonies. **B**. Small and large colonies isolated under conditions described above were patched on YP+2% glycerol (YPG) medium and incubated at 30°C for 2 days.

Petite mutants are resistant to FLU, primarily through up-regulation of efflux pumps (49). We, therefore, repeated FIC assays, combining FLU with 4377 in YPD instead of the standard RPMI media to facilitate growth of petite strains; we also carried out the FIC in YPG to suppress the formation of respiratory-deficient strains. In addition, we measured cell density rather than using a visual readout to allow for a more nuanced assessment of the activity of the combinations. Both 4377 and FLU were more modestly more active in YPG as compared to YPD (**Fig. 9A**); to our knowledge, this effect of glycerol medium on FLU has not been reported previously. Glycerol medium also potentiated the activity of the combination as evidenced by smaller area of minimal inhibition for the YPG experiment; the same effect was observed for all other MEF-derivatives tested. The contents of combination wells with approximately 50% inhibition were plated on YPD and incubated at 30°C. The majority of cells isolated from these plates were morphologically petite (**Fig. 9B**) and 20/20 representative petite colonies were unable to grow on glycerol media (data not shown). DAPI staining of the respiratory deficient cells showed decreased mitochondrial DNA staining, confirming these strains had alterations in the DNA content of their mitochondria (**Fig. 9C**). Thus, the ability of MEF-derivatives induce petite formation which, in turn, appears to modulate the activity of FLU/MEF analog combinations. Importantly, we did not observe this phenomenon with *C. neoformans* or *C. albicans*, most likely because these strains are so-called “petite-negative” (data not shown) and cannot grow if their mitochondrial DNA has been lost (50). Taken together, these experiments strongly support the notion that MEF-derivatives interfere with mitochondrial function but, like the vacuolar effects, the mitochondrion is not be the sole target driving antifungal activity. Rather, the activity of MEF and its derivatives is best explained by a combination of effects on the mitochondrial and vacuole.

**Figure 9.**
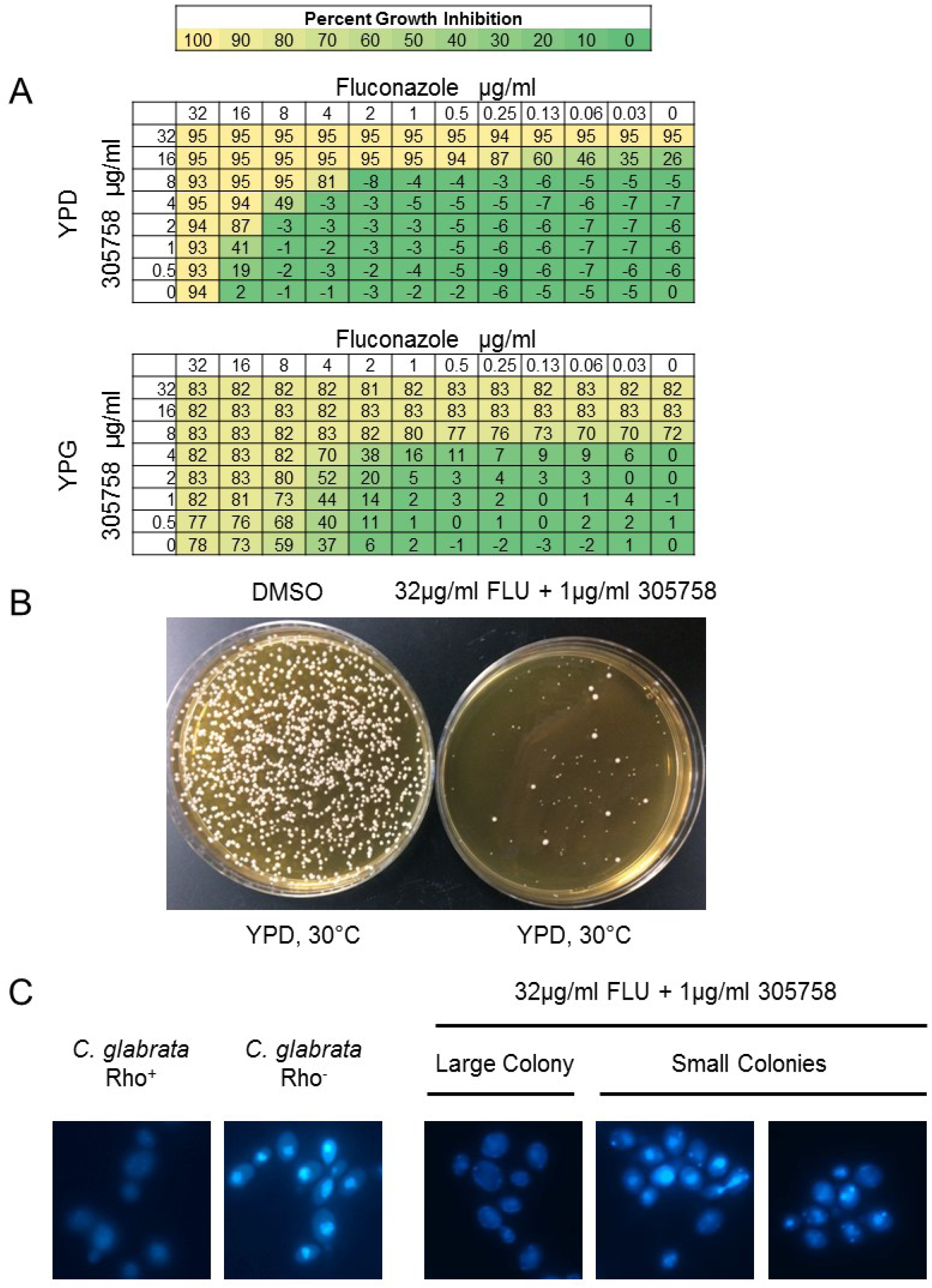
*C. glabrata* petite formation modulates the activity of mefloquine-derivative/fluconazole combinations. **A.** Checkerboard assays with *C. glabrata* BG2 were performed with 4377 and fluconazole in YPD and YPG at 37°C. After 24 hr incubation, the optical density of each well was measured and compared to the DMSO only well. The percentage of growth inhibition was calculated and is shown both numerically and in heat map form. **B.** The surviving cells from a well with approximately 50% growth inhibition as well as the DMSO only well were plated on YPD and incubated for 2 days at 30°C. **C.** *C. glabrata* Rho+ strain KK2001 and a Rho-derivative were stained with DAPI. The Rho+ strains show a diffuse staining pattern indicative of both mitochondrial and nuclear DNA staining while Rho-strains show a focal nuclear pattern indicative of nuclear staining only. Large colony cells stain in a pattern consistent with Rho+ while the petite colony cells show the focal pattern consistent with Rho^−^.

## DISCUSSION

The small number of drugs and drug classes that can be used to treat invasive, life-threatening fungal infections necessitates an expansion of the types of approaches used to find new drugs as well as the types of molecular classes that are considered for further development. Drug repurposing has emerged as an attractive approach to expediting the drug development process. In general, such repurposing efforts have focused on identifying a specific drug that can be directly transitioned to a new clinical use without modification of the structure or formulation. Here, we adopted a repurposing approach toward a different goal: to identify a currently used drug with antifungal properties as a point of departure for further development and optimization.

One of the most compelling reasons to pursue repurposing approaches to identifying new drug candidates is that the molecules and scaffolds have known pharmacology. The pharmacologic properties of a new chemical entity are very difficult to predict, a priori and it is frequently difficult to redress serious metabolic or toxicologic liabilities by re-engineering the scaffold. Accordingly, we chose to explore the antifungal properties of MEF further is that its pharmacology has a number of useful features for the treatment of fungal infections (51). First, it is well absorbed and establishes high serum and tissue levels when taken orally. Second, it penetrates the blood-brain-barrier and achieves high concentrations relative to the plasma (52). Although new therapies for the neurotropic pathogen *C. neoformans* represent one of the most important unmet clinical needs in medical mycology (25), it is important to recognize that almost all invasive fungal infections can lead to CNS disease. Third, the use of fungal prophylaxis of high-risk patients has become established practice for many pre-disposing conditions (3). MEF has a long half-life and, thus, could provide dosing regimens that would be quite amenable to prophylaxis. It is important to note that MEF has well-documented neurological/psychiatric side effects in a minority of patients (53). These are generally associated with long term use in the setting of malaria prophylaxis. If molecules of this class ultimately progress through pre-clinical development, then it will be important to carefully consider the risk-benefit ratio of their use. However, the *in vitro* activity and pharmacologic properties of the class seem worthy of consideration at this time.

The MEF scaffold had been known to have antifungal properties for over twenty years so our project represents something of a rediscovery (22); however, no in-depth characterization of the antifungal properties of MEF or its derivatives has been undertaken. As other’s have shown MEF has poor to modest *in vitro* antifungal activity against the *Candida* spp., *Cryptococcus*, and *Aspergillus*. In contrast, the MEF-derivatives that we obtained from the NCI depository are much more active (up to 32-fold more active) against a broad spectrum of pathogens including *Candida* spp., *C. neoformans*, and *A. fumigatus*. Overall, NCI 4377 is the most active analog with MICs between 1 and 4 µg/mL for the species tested but, all four of the most active derivatives had MICs in the 2-8 µg/mL against the species tested.

Despite the significant increase in antifungal activity relative to MEF, the *in vitro* toxicity of the MEF-derivatives against human cells was only minimally increased. This differential increase in antifungal activity is particularly important because it indicates that the MEF-derivatives are not simply more active generalized eukaryotic poisons. Indeed, Dow et al. found that both NCI 2450 and NCI 305758 are less neurotoxic than the parent MEF derivative (52). A large number of MEF-derivatives and alkylamino-quinolines in general have been synthesized and evaluated as anti-malarial molecules which might be exploited for an expedient SAR-based optimization of the antifungal activity of this scaffold (54).

The recent emergence of drug-resistant and multi-drug resistant fungi has generated well-founded concerns in the face of the limited antifungal repertoire (55). Thus, the activity of the MEF-derivatives against multi-drug resistant *C. auris* and *C. glabrata*, two of the most important drug-resistant organisms, is very promising. This feature indicates that the class is not susceptible to the mechanisms of resistance such as increased efflux pump expression that affect azole drugs. These observations also suggest that the mechanism and/or mode of action of the MEF class is distinct from antifungal drugs currently in use. Based on these characteristics, further medicinal chemistry optimization of the antifungal activity appears to be warranted.

The ability of MEF and its derivatives to achieve high concentrations in the CNS makes them attractive candidates for use in the treatment of cryptococcal meningitis (52). Currently, the gold standard therapy for cryptococcal meningitis is AmB combined with 5-flucytosine. As has been well-documented, this regimen is not available in many resource-limited regions of the world with high burden of cryptococcal meningitis (26). FLU is far less effective because of its fungistatic mode of action but is much easier to obtain and administer. The development of adjuvants to, or combinations with, FLU have been of interest to the field. Ideally, an orally available drug could be combined with FLU to yield a fungicidal cocktail with clinical efficacy similar to the AmB and 5-flucytosine combination. Our *in vitro* time-kill studies indicate that MEF and its derivatives are fungicidal when combined with FLU and, thus, are candidates for combination therapy.

The exact mechanism of action for the anti-malarial activity of MEF has remained ill-defined despite its longstanding use. Although our studies on its mechanism of action as an antifungal have not identified a single protein target, new insights into the cellular effects and mode/mechanism of action have emerged. First, our data are most consistent with the idea that the antifungal activity of MEF and its derivatives is due to effects on more than one target. This conclusion is supported by the fact that we could not isolate stable resistant-mutants by *in vitro* selection.

Second, we propose that at least part of the antifungal activity of the MEF class is due to its ability to interfere with the function the function of both the mitochondria and the vacuole. Other antifungal molecules that appear to more specifically target the mitochondria have been identified; however, these molecules typically show increased activity in the presence of non-fermentable carbon sources such as glycerol relative to fermentable carbon sources such as glucose. The mefloquine derivatives do not show this effect despite disrupting the proton motive force across the mitochondrial membrane and interfering with mitochondrial DNA stability. Thus, the mitochondria cannot be the sole target. In keeping with our first conclusion, other effects such as those manifested by vacuolar disruption or secretory pathway blockade are likely to combine with mitochondrial disruption. This proposal is also supported by previous chemical-genetics data indicating that strains with mutations in vacuolar and secretory proteins are hypersusceptible to MEF. Finally, we also assert that the lack of a defined, specific target explaining the anti-malarial activity of MEF may be because, as with its antifungal activity, it has multiple effects and targets.

In summary, the broad-spectrum activity, known and favorable pharmacological properties, and the novel, multi-target mechanism of action strongly support the development and optimization of the antifungal activity of the MEF scaffold.

### Yeast strains and general microbiological methods

Media was prepared using standard recipes (56). All fungal strain glycerol stocks are stored at −80°C and were propagated on yeast peptone dextrose (YPD) agar plates (1% (wt/vol) yeast extract, 2% (wt/vol) peptone, 2% (wt/vol) dextrose and 2% (wt/vol) agar. All strains were used within 10 days of streaking. *C. albicans* strain SC5314 was obtained from Gus Haidaris (University of Rochester). *C. albicans* and *C. glabrata* clinical isolates were obtained from Ellen Press (University of Pittsburgh). *C. auris* clinical isolates were obtained from Daniel Diekema (University of Iowa). *C. neoformans* clinical isolates were obtained from John Perfect (Duke University). Unless otherwise indicated, experiments began with an initial culture of cells that were grown overnight (∼16 hr) in YPD liquid media, shaking at 30°C. All drugs were solubilized in DMSO.

### *In vitro* antifungal susceptibility

Antifungal susceptibility testing was performed using the protocols described in the document CLSI M27-A3. The reported minimum inhibitory concentration (MIC) values represent the highest value for at least three independent biological replicates performed in at least technical duplicate. Checkerboard fractional inhibitory concentration (FIC) assays were performed as previously described using the same medium and conditions as the CLSI M27-A3 protocol (57). MIC and FIC results were read by visual inspection and/or by cell optical density readings at absorbance 600nm (OD_600_).

### Time-kill assay

Overnight cultures of *C. neoformans* (H99) were diluted in fresh YPD to an OD_600_ of 0.1-0.2 and grown at 30°C shaking until OD_600_ = ∼ 0.4. Once OD_600_ was reached, cultures were incubated at 37°C shaking with vehicle or the indicated concentration of drug (final DMSO concentration was 0.4%) for 24h. The 0h time point was taken before the addition of drug and subsequent samples were taken every 4h after the addition of drug. At each time point, serial dilutions of samples were plated on YPD agar plates, incubated at 30°C, and counted after 48h. Experiments were done with at least two independent biological replicates and in technical duplicates.

### *In vitro* toxicity assays

Cell lines were obtained from ATCC and included A549, human alveolar cells and HepG2, human liver cells. Cells were cultured with 10% FBS, 1% Penicillin/Streptomycin, and 1% glutaMAX in DMEM/F-12 15mM HEPES (A549) or DMEM (HepG2). Toxicity and viability were measured using an XTT reduction assay. Cells were seeded and incubated at 37°C in 5% CO_2_ until cells were ∼70% confluent (24 – 48hr). Media was aspirated and replenished in combination with DMSO or serial dilutions of drug. Cells were incubated in drug or DMSO under the same conditions for an additional 24h. Assay kits, Pierce LDH Cytotoxicity Assay Kit (Catalog # 88954) and CyQUANT™ XTT Cell Viability Assay (Catalog # X12223) were used, developed, and measured according to each assay protocol.

### *C. albicans* filamentation

Overnight cultures of *C. albicans* (SC5314) were diluted 1:100 and incubated with drug or DMSO (0.5% DMSO in final culture) in YPD for 1h shaking at 37°C. After 1h, cultures were induced with 1%FBS and incubated for an additional hour shaking at 37°C. Samples were centrifuged, resuspended in YPD, stained with propidium iodide 10µg/ml, incubated in the dark for 30 minutes, washed once with DPBS, resuspended with DPBS and imaged. Images were captured using a Nikon ES80 epifluorescence microscope and visualized with a CoolSnap CCD camera and NIS-Elements Software. Quantitative analysis was performed by visually inspecting images and hyphae were measured using the NIS-Elements Software. The data represent the mean of at least two biological replicates for which at least 100 cells were counted. Error bars indicate standard deviation.

### *C. neoformans* melanization and capsule formation

Overnight cultures of *C. neoformans* (H99) were washed in DPBS and resuspended to OD_600_= 1.0. 1:10 dilutions were made in sterile UltraPure™ DNase/RNase-Free Distilled Water. Of those dilutions, 10 µl was spotted onto L-DOPA plates that contained either DMSO or drug in duplicate with at least two biological replicates. Plates were incubated at 37°C for 48h and imaged.

### Vacuole morphology

Cells were grown overnight in YPD, shaking at 30°C. Cells were back diluted fresh YPD to an OD_600_=0.2-0.3 and grown at 30°C until cells reached log phase growth (OD_600_=0.5-0.7). Cells were then exposed to DMSO or drug and incubated at 30°C for 2h shaking. After 2h, samples were centrifuged, resuspended in YPD with 40µM FM4-64, and incubated in the dark shaking at 30°C for 15 minutes. Cells were centrifuged, reuspended in YPD in a dark tube, and incubated shaking at 30°C for 1h. Cells were then centrifuged, resuspended in DPBS, mounted and imaged on the aforementioned epifluorescent microscope.

### Mitochondrial assays

Cells were grown overnight in YPD, shaking at 30°C. The next day overnight culture was back diluted into fresh YPD to an OD_600_=0.2-0.3 and grown at 30°C shaking until the OD_600_=0.5-0.7. Once OD_600_ was reached, Cells were exposed to DMSO or drug for 2h for *C. albicans* (SC5314) or was exposed for 30 minutes for *C. neoformans* (H99) at 30°C shaking. MitoTracker™ Red CMXRos (Catalog # M7512) and/or MitoTracker™ Green FM (Catalog #M7514) was prepared in warm YPD for a final concentration of 1mM. Once drug incubation was complete, cells were centrifuged and resuspended with the YPD and MitoTracker mixture, then incubated in the dark at 30°C for 50 minutes. Cells were centrifuged, washed with fresh YPD, resuspended with PBS, mounted to a slide, and imaged.

### Identification and Validation of Petites

MICs were done as previously described with wild type *C. glabrata* KK2001 and plates were read at 24h and 48h. At 48h, the wells that grew past the recorded 24h MIC and control wells were diluted, spotted and plated on YPD and YPG plates. Plates were incubated at 30°C overnight and the next day, petites and larger control colonies from the YPD plates were patched onto fresh YPG plates, incubated at 30°C overnight to confirm petite phenotype. Glycerol stocks were made for future DAPI staining experiments. The same procedure was done for FIC plates of mefloquine analog and fluconazole. Petites were verified via a secondary assay of DAPI staining. Briefly, petite and control glycerol stocks were streaked on YPD plates and grown for 48h at 30°C. Overnight cultures were grown at 30°C for 16h and the following day, cells were washed in synthetic complete media, back diluted in synthetic complete media to an OD_600_ = 0.2-0.3 and grown to log phase at 30°C shaking until the OD_600_= 0.4-0.6. Once OD was reached, cells were stained with 2.5µg/ml DAPI and grown for 30 minutes at 30°C shaking. Cells were then centrifuged, resuspended in fixing reagent containing 4% formaldehyde in synthetic complete media, and incubated at room temperature for 15 minutes. Then cells were washed with DPBS, resuspended in DPBS, mounted on slides, and imaged on the epifluorescent microscope. Cells were always imaged the same day as fixing, not after storage.

---

**Table S1.**
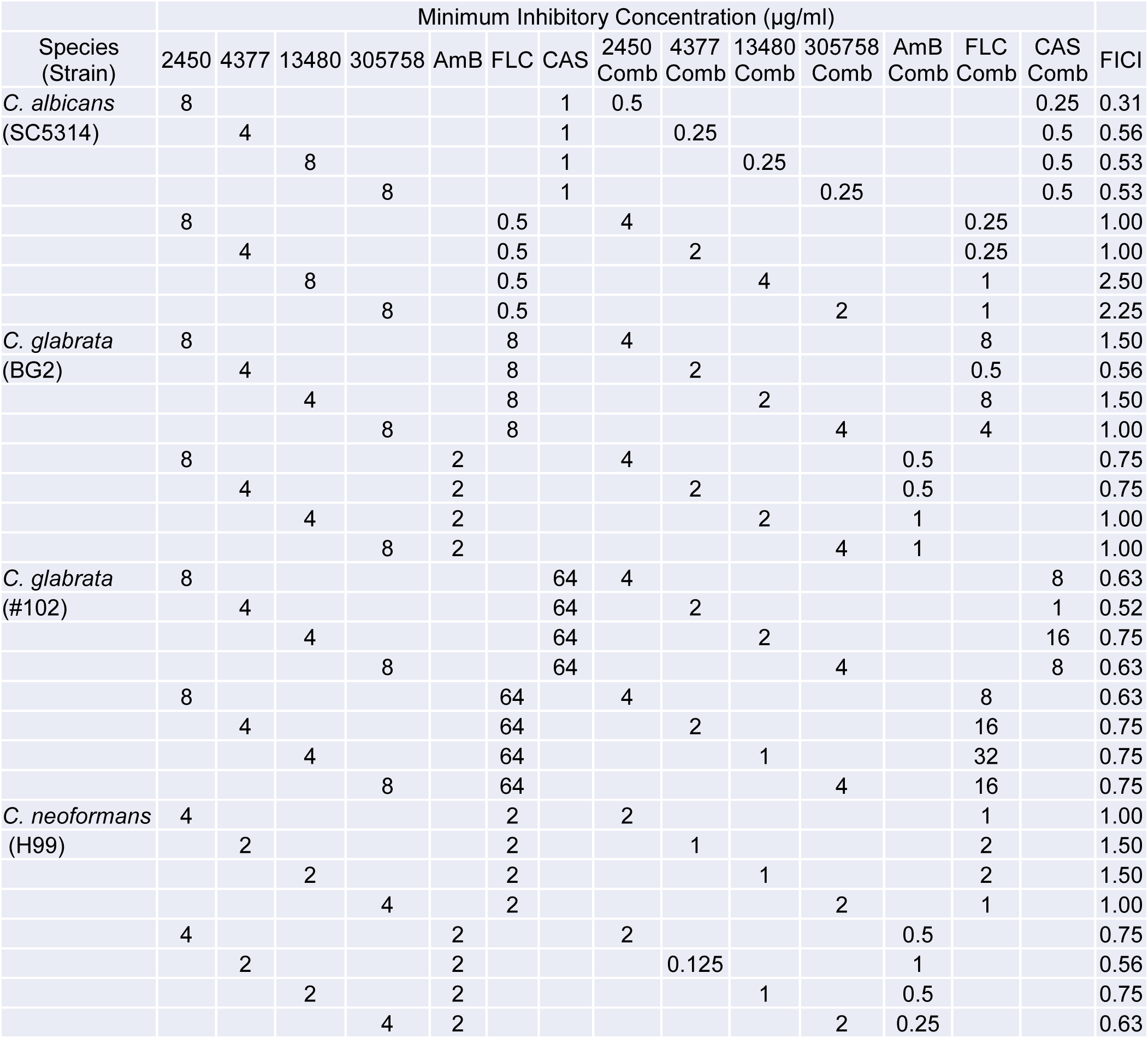
Fractional Inhibitory Concentrations of Mefloquine Derivatives with Clinical Antifungals against Susceptible Strains of C. albicans, *C. glabrata*, and *C. neoformans* or a Resistant Clinical Isolate of *C. glabrata* (#102).

